# Spinal Recurrent Inhibition Shapes the Dynamics of TMS-induced Motor-Evoked Potentials

**DOI:** 10.64898/2026.09.11.750929

**Authors:** Vincent S.C. Chien, Eleonora Bernasconi, Erik Müller, Peng Wang, Madeleine Lowery, Jérémy Liegey, Karen Wendt, Jacinta O’Shea, Timothy Denison, Federico Ranieri, Vincenzo Di Lazzaro, Jaroslav Hlinka, Thomas R. Knösche, Konstantin Weise, Helmut Schmidt

## Abstract

**Background:** Motor-evoked potentials (MEPs) elicited by TMS reflect the integrity of the corticospinal pathway, yet the contributions of spinal and peripheral mechanisms to the shape of individual MEP waveforms remain poorly understood.

**Objective:** To develop a biologically grounded model of the spinal and peripheral motor pathway that can be fitted to individual MEP waveforms across TMS intensities, and to test whether Renshaw cell (RC) recurrent inhibition is required to reproduce them.

**Methods:** Corticospinal volleys (D- and I-waves), reconstructed from published epidural recordings in humans, drove 100 conductance-based leaky integrate-and-fire alpha motor neurons coupled to an RC population. Motor unit action potentials were derived from a finite-element hand model informed by diffusion tensor imaging of the first dorsal interosseous muscle. Models with (RC^+^) and without (RC^*−*^) recurrent inhibition were fitted to MEPs across TMS intensities in 10 healthy participants and compared with a phenomenological benchmark.

**Results:** The RC^+^ model explained approximately 90% of waveform variance (mean *R*^2^ = 0.89), comparable to the benchmark (0.91; *p* = 0.32). Removing RC inhibition reduced waveform goodness-of-fit in every participant (mean *R*^2^ = 0.79; *p* = 0.002 vs. RC^+^), although input-output curve fits of MEP amplitude did not differ significantly between the two variants (*p* = 0.49).

**Conclusions:** RC recurrent inhibition was required to reproduce the temporal structure of TMS-evoked MEPs. MEP waveforms thus carry more information than amplitude-based measures. Fitted to individual waveforms, the model can estimate unobservable spinal and peripheral activity, such as motor unit firing times, as well as participant-specific pathway parameters.

## 1 Introduction

The motor-evoked potential (MEP) elicited by transcranial magnetic stimulation (TMS) and recorded from peripheral muscles using surface electromyography (EMG) reflects cortical excitability and the integrity of the corticospinal tract [1, 2]. MEP recordings play an important role in the prognosis of motor recovery after stroke [3, 4], the diagnosis of various neurological disorders [5, 6], the evaluation of spinal cord injury in clinical trials [7], and intraoperative monitoring [8].

A TMS pulse over the primary motor cortex produces descending volleys in the corticospinal tract, consisting of a direct (D-) wave arising from direct activation of corticospinal axons and indirect (I-) waves generated by trans-synaptic activation of cortical interneurons (collectively, DI-waves) [2, 9]. In the spinal cord, these volleys are integrated by alpha motor neurons (*α*MNs) and modulated by local inhibitory interneurons such as Renshaw cells (RCs). The descending signals ultimately drive the contraction of innervated muscles. Muscle activity generates motor unit action potentials (MUAPs), which summate to form the MEP recorded in surface EMG.

Previous computational studies have provided insights into the cortical generation of TMS-induced DI-waves [10, 11], and some of these studies included peripheral models to describe MEP generation [12, 13], targeting the input-output (IO) curve (i.e., MEP peak-to-peak amplitude against TMS intensity) and the facilitation and inhibition induced by paired-pulse protocols. However, these models restricted the spinal component to MN populations alone, omitting inhibitory interneurons such as RCs, and used simplified representations of MUAPs, focusing on MEP peak-to-peak amplitude rather than MEP’s fine temporal structure. A parallel line of work has developed biophysically detailed models of spinal MN pools with realistic EMG and force output [14–17]. Although some include RCs, they are oriented toward proprioceptive feedback and voluntary motor control rather than TMS-evoked responses. RC recurrent inhibition is known to regulate MN synchronization and the temporal structure of motor output, acting as a neural filter relevant to tremor and fine motor control [18, 19]. Whether it also shapes the temporal structure of TMS-evoked MEP waveforms, a transient, open-loop context distinct from voluntary contraction, has not been examined.

Moreover, existing MEP models have accounted for group-level phenomena but have not been fitted to individual participants. Individual MEP waveforms vary not only with recording conditions but also with individual differences along the motor pathway. A model that is flexible enough to capture individual characteristics, yet computationally efficient and biologically plausible enough for its fitted parameters to be interpretable, could help disentangle the contributions of cortical, spinal, and peripheral stages to the observed MEP.

To address these needs, we developed a spinal-peripheral model (Figure 1) designed to fit individual MEP waveforms across a range of TMS intensities. D- and I-waves, reconstructed for each TMS intensity from group-level DI-wave IO curves measured with epidural recordings in humans [20], serve as model input and drive 100 conductance-based leaky integrate-and-fire (LIF) *α*MNs that interact with a common inhibitory RC pool. Each MN and the muscle fibers it innervates form a motor unit (MU), whose MUAP is derived from a finite-element (FE) volume-conductor model of the hand informed by diffusion tensor imaging (DTI) of the first dorsal interosseous (FDI) muscle, and the MEP is computed as the sum of time-shifted MUAPs. We fitted model variants with (RC^+^) and without (RC^*−*^) recurrent inhibition to MEPs from 10 healthy participants [21] and compared them with a phenomenological benchmark developed in this study, which bypasses the spinal circuit and samples MU firing times directly from a gamma distribution, in terms of the goodness-of-fit (GoF) of both MEP waveforms and IO curves.

**Figure 1.**
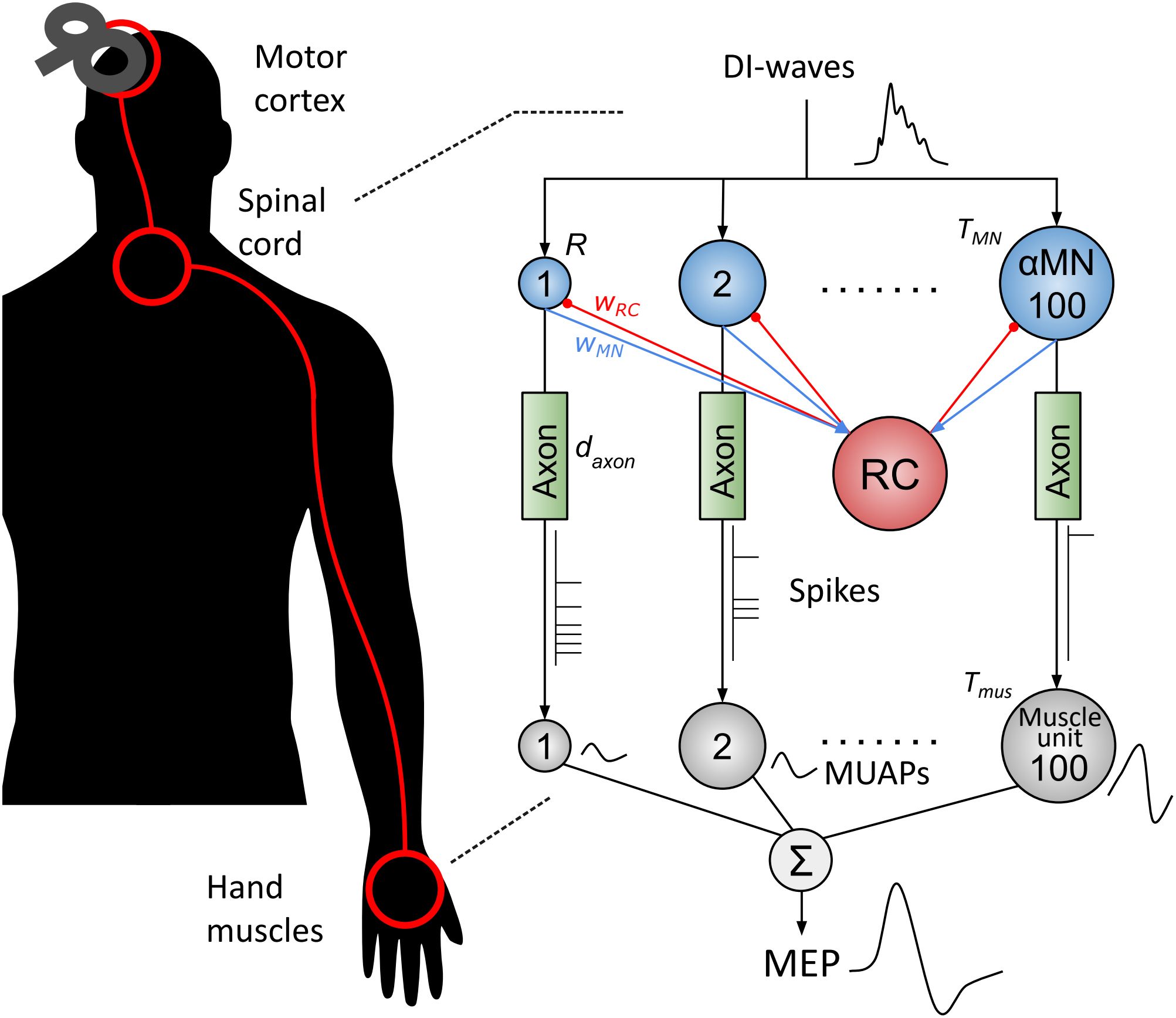
The structure of the spinal-peripheral model. (Left) TMS over the motor cortex evokes descending D- and I-waves (DI-waves) in the corticospinal tract, which drive spinal motor neurons whose axons project to the hand muscles. (Right) The model comprises a spinal cord component and a hand muscle component. The spinal cord component consists of 100 alpha motor neurons (*α*MNs, blue circles, indexed 1 to 100 and sorted by membrane resistance *R* in descending order) receiving the same DI-wave input, and a single Renshaw cell (RC) population. The *α*MNs excite the RC population with connection weights *w*_MN_ (blue lines), and the RC population feeds inhibition back to all *α*MNs with connection weights *w*_RC_ (red lines with round ends). Each *α*MN has a fixed refractory period *T*_MN_. The hand muscle component consists of 100 muscle units (gray circles), each innervated by its corresponding *α*MN via a motor axon with a common axonal conduction delay *d*_axon_. Each motor unit (MU), consisting of an *α*MN and its muscle unit, generates a motor unit action potential (MUAP, waveforms next to the gray circles) whenever an MN spike (spike trains below the axons) arrives outside the muscle unit refractory period *T*_mus_. The MEP is computed as the summation (Σ) of time-shifted MUAPs across all MUs, as recorded by surface EMG.

## 2 Methods

### 2.1 MEP dataset

The MEP dataset of 10 healthy participants was recorded and published in [21] and shared via the Open Science Framework (OSF, https://osf.io/5ry92/). The original data collection protocol was approved by the Central University Research Ethics Committee (CUREC), University of Oxford (R75180/RE002), and all participants provided written informed consent prior to participation [21]. The recording procedure is described in detail in [21] and summarized below. The TMS stimuli were applied in blocks of 15 in increasing order from low to high intensities in steps of 3% of the maximum stimulator output (MSO) using a Magstim 200^2^ stimulator and a focal 70-mm figure-of-eight coil. The coil was held over the left primary motor cortex at the hotspot of the right first dorsal interosseous (FDI) muscle, oriented at 45°to the midline with the handle pointing posteriorly to induce a posterior-anterior (PA) current. Within each block, the inter-pulse intervals were varied between 4.25 and 5.75 s. EMG was recorded at rest from the first dorsal interosseous muscle of the right hand by positioning disposable neonatal ECG electrodes in a belly-tendon montage, with the ground electrode over the ulnar styloid process. The EMG signals were recorded using a D440 Isolated Amplifier (Digitimer, Welwyn Garden City, UK), a Micro1401 (Cambridge Electronic Design, Cambridge, UK), a Digitimer HumBug Noise Eliminator and Signal version 7.01 (Cambridge Electronic Design), with a 16-bit resolution at a 10 kHz sampling rate, an amplifier gain of 1000 and a 10–1000 Hz filter.

### 2.2 Spinal-peripheral model

The model comprises a spinal cord component and a hand muscle component (Figure 1). It receives descending DI-waves as input and generates MEP waveforms as output.

#### 2.2.1 DI-wave input

The model input consists of DI-waves, which reflect the direct (D-wave) and indirect (I-wave) corticospinal volleys elicited by TMS in the motor cortex. Because participant-specific DI-wave recordings were not available, we reconstructed DI-waves from group-level DI-wave IO curves extracted from Figure 2 of [20], which reports the amplitudes of D- and I-waves recorded epidurally from the high cervical cord at rest in patients with chronic back pain (Figure 2A). These volleys were evoked by a figure-of-eight coil held over the right motor cortex at the optimal scalp position for eliciting motor responses in the contralateral FDI, with the induced current flowing in a PA direction. These stimulation parameters match those of the MEP dataset analyzed here (Section 2.1, MEP dataset), except for the stimulated hemisphere. For each participant, TMS intensities (%MSO) were expressed relative to the individual resting motor threshold (RMT) and mapped onto these IO curves. For each TMS intensity, the DI-waves were modeled as five Gaussian components with peak amplitudes given by these IO curves (Figure 2B, top). The D-wave was placed at 2.5 ms, consistent with D-wave latencies recorded epidurally from the high cervical cord [9, 22], and the I-waves followed at intervals of 1.5 ms (I1–I4 at 4 ms, 5.5 ms, 7 ms and 8.5 ms), corresponding to the typical periodicity of I-waves [9, 23]. Each component had a width of 0.25 ms to account for conduction delay variability along corticospinal axons [24]. The DI-waves were then deconvolved with an extracellular potential (EP) kernel (Figure 2B, middle) to estimate the underlying DI-wave firing rates *m*_DI_(*t*), which serve as model input (Figure 2B, bottom). The computation of the EP kernel is described in the Supplementary Material.

**Figure 2.**
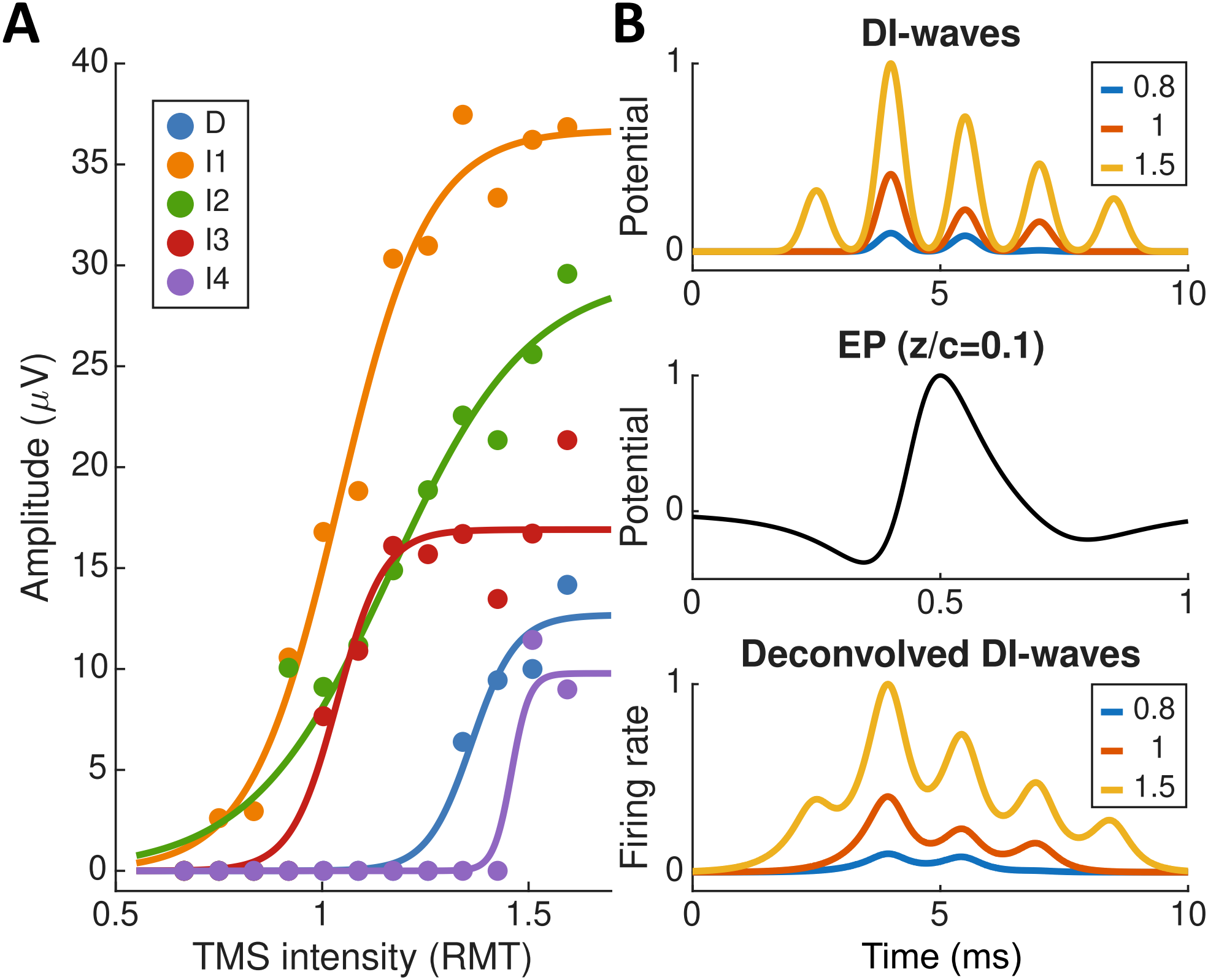
Reconstructed DI-wave input. **(A)** DI-wave input-output (IO) curves. The colored dots, digitized from the experimental observations reported in Figure 2 of [20], describe the average peak-to-peak amplitudes of the epidural D- and I-waves (I1–4) elicited by different TMS intensities (relative to the resting motor threshold, RMT). The DI-wave IO curves are derived by fitting five sigmoid functions to the dots. **(B)** Examples of reconstructed DI-waves at 0.8, 1, and 1.5 RMT intensities (top row). The reconstructed DI-waves consist of five Gaussian peaks, with peak amplitudes corresponding to the amplitudes at a given TMS intensity in the IO curves in (A). The five Gaussian peaks have a width of 0.25 ms and a peak-to-peak interval of 1.5 ms [9, 23, 24]. The reconstructed DI-waves are deconvolved with an extracellular potential (EP) kernel (middle row) to yield deconvolved DI-waves (bottom row), which are estimated spike volleys in the spinal cord and serve as model input.

#### 2.2.2 Spinal cord component

The spinal cord component considers the interaction between *N*_MU_ = 100 conductance-based LIF MNs (where *N*_MU_ denotes the total number of motor units) and a single RC population. It receives descending DI-waves from the corticospinal tract and generates MN spikes that are transmitted through motor axons to activate the hand muscles.

Each MN is modeled as an LIF neuron with membrane time constant *τ* = 10 ms, resting potential *V*_rest_ = −65 mV, and firing threshold *V*_th_ = −55 mV. After each spike, the membrane potential is reset to *V*_rest_ for a refractory period of *T*_MN_ = 2 ms. The dynamics of the *i*-th MN are described by

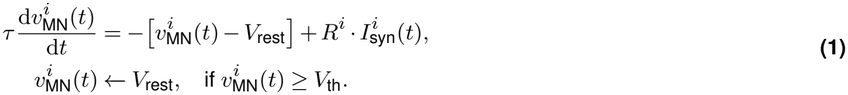

The synaptic current comprises an excitatory component driven by the DI-wave input and an inhibitory component mediated by RC feedback:

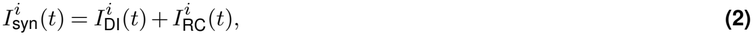

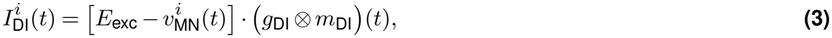

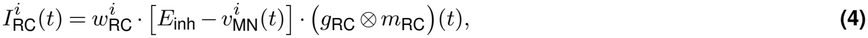

where *E*_exc_ = 0 mV and *E*_inh_ = −75 mV are the reversal potentials, *⊗* denotes convolution, *g*_DI_(*t*) and *g*_RC_(*t*) are the synaptic kernels for the excitatory and inhibitory projections, respectively, *m*_RC_(*t*) is the firing rate of the RC population, and 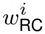 is the RC*→*MN weight on the *i*-th MN. The synaptic kernels combine AMPA and NMDA receptor-mediated components for the excitatory Cortex*→*MN projection and a glycinergic component for the inhibitory RC*→*MN projection:

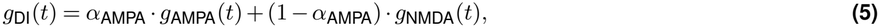

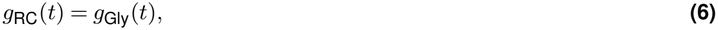

where *α*_AMPA_ *∈* [0, 1] is a free parameter. Each basis kernel follows a biexponential form,

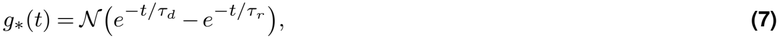

with rise and decay time constants *τ*_*r*_ and *τ*_*d*_ listed in Table 1. The normalization *N* (*·*) scales the peak conductance to 0.01 nS, shifting receptor-specific differences in peak conductance to the free parameters *α*_AMPA_, *R*, and *w*_RC_.

**Table 1.** Parameters of synaptic kernels.

| Synapse | τ<sub>r</sub><br>(ms) | τ<sub>d</sub><br>(ms) | Ref. |
|---|---|---|---|
| Cortex→MN (AMPA) | 1 | 5 | [25] |
| Cortex→MN (NMDA) | 3 | 50 | [25] |
| RC→MN (Glycine) | 1 | 6 | [26, 27] |
| MN→RC (fast ACh) | 0.5 | 3.6 | [28] |
| MN→RC (slow ACh) | 1.8 | 20.2 | [28] |

RCs receive excitatory input from MN axon collaterals and inhibit the same and neighboring MNs, thereby preventing excessive MN firing [29–32]. To reduce model complexity, a single RC population represents the average RC response. The MN spikes 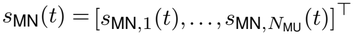, where *s*_MN,*i*_(*t*) = ∑_*n*_ *δ t − t*_*i,n*_ and *t*_*i,n*_ are the motor neuron spike times, reach the RC population with connection strengths 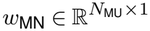 and induce excitatory postsynaptic potentials (EPSPs) *v*_*\**_(*t*) that follow a second-order equation implementing the biexponential kernel (Eq. 7):

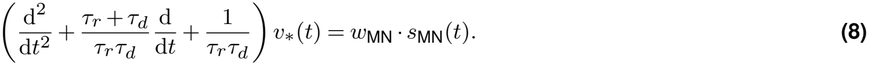

Fast and slow acetylcholine (ACh) receptor components [28] are combined with a relative weight *α*_fACh_ = 0.5,

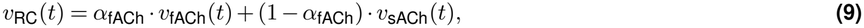

and the RC firing rate *m*_RC_(*t*), constrained to [0, 1] Hz, is obtained through a sigmoid with slope *r* = 10 mV^−1^ (fixed during parameter selection; see Discussion) and threshold *v*_0_ (a free parameter ranging from 1 to 10 mV):

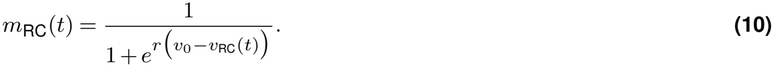

#### 2.2.3 Hand muscle component

A motor unit (MU) consists of a single MN and all of the muscle fibers it innervates (its muscle unit). When the MN generates an action potential, the signal propagates along its axon and activates all innervated muscle fibers. The summed extracellular potential generated by these fibers during a single firing of the MU is known as a MUAP.

A repertoire of MUAP waveforms *u*_*i*_(*t*), *i* = 1, …, *N*_MU_, one for each MN, was simulated according to [33], using an FE model of the hand derived from anatomical MRI data (Figure 3A,B), together with DTI fiber tracking. The FE model was adapted to incorporate a 28 mm × 20 mm monopolar EMG electrode (Ambu Neuroline 710) at the skin surface. The trajectories of individual muscle fibers within the FDI muscle were estimated based on the DTI tracts (Figure 3C). Muscle fibers were randomly distributed throughout the muscle cross-section and assigned to MUs, with overlapping territories according to MU size as defined in [34] (Figure 3D). For each MU, the extracellular potential generated by its constituent fibers at the recording electrode was calculated by solving the Laplace equation using the FE model and making use of the principle of reciprocity. The muscle fiber action potentials of each MU were summed to yield the MUAP waveform *u*_*i*_(*t*) corresponding to each MN in the spinal cord component (Figure 3E).

**Figure 3.**
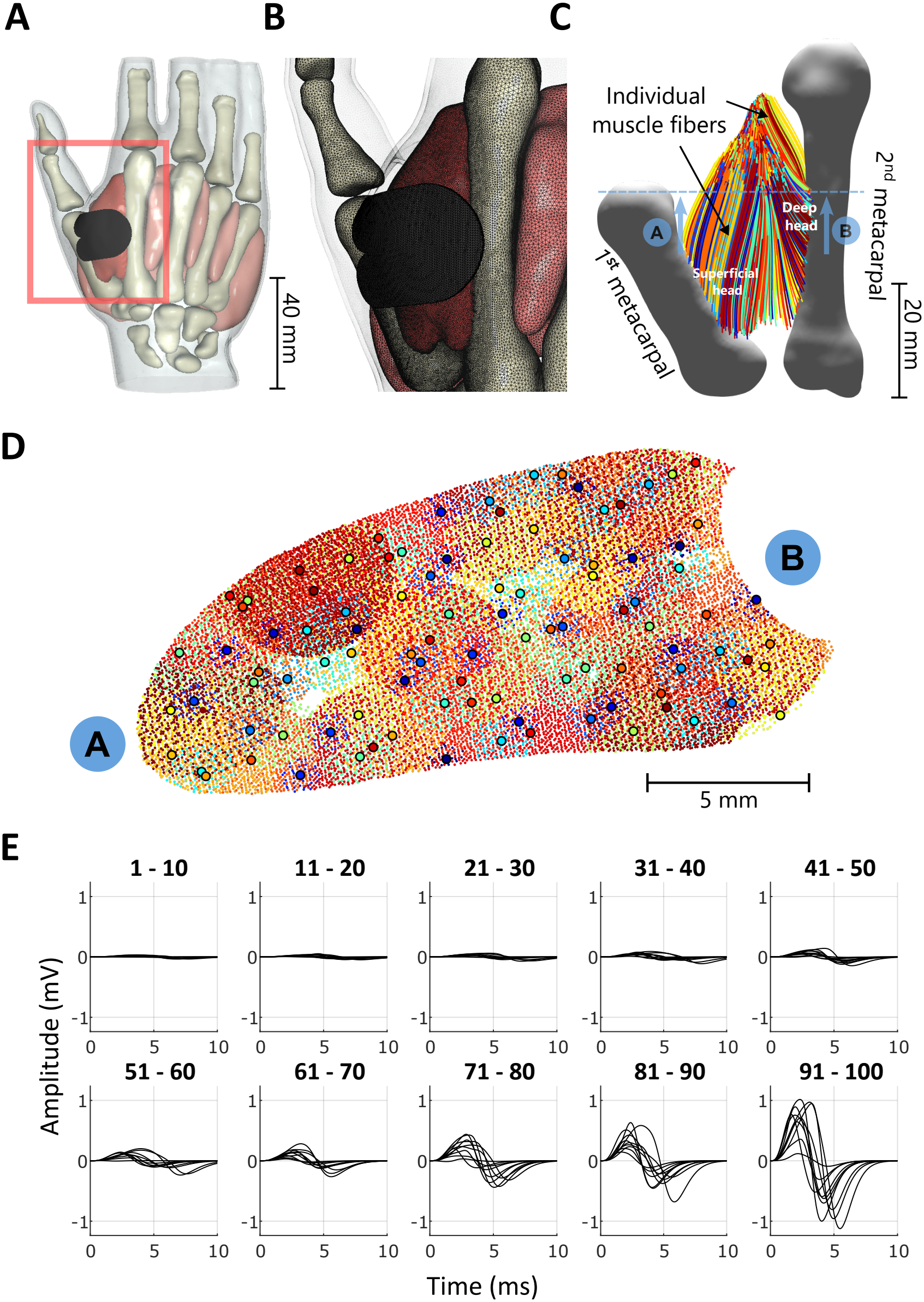
Anatomically derived MUAP repertoire. **(A)** Segmented hand model derived from MRI data, including the metacarpal bones and hand muscles (FDI shown in red), and surface electrode (dark gray) highlighted by the rectangle. **(B)** Finite-element discretization (tetrahedral mesh) of the hand model shown in (A). **(C)** Dorsal view of the reconstructed FDI muscle fiber trajectories (colored lines), derived from diffusion tensor imaging (DTI) fiber tracking, spanning the 1st and 2nd metacarpal bones; the dashed line indicates the cross-section shown in (D). **(D)** FDI cross-section directly beneath the surface electrode. Colored dots indicate individual muscle fibers, and colored discs mark the centers of the corresponding motor unit (MU) territories. **(E)** Repertoire of *N*_MU_ = 100 MUAPs simulated following [33]. The number above each subplot indicates the range of MU indices shown. The waveforms were normalized so that the maximum MUAP amplitude across the MU population equals one. The MUAP index corresponds to the MN index.

MN spikes reach the muscle unit after a common axonal delay *d*_axon_, and each MN spike that arrives outside the muscle unit refractory period *T*_mus_ generates a MUAP. The resulting MU firing times 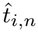, i.e., the times at which the *i*-th MU generates its *n*-th MUAP, define the MU spike train:

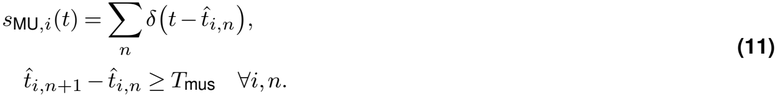

MU firing times thus form a subset of the delayed MN spike times. The simulated MEP waveform is the sum of all time-shifted MUAPs:

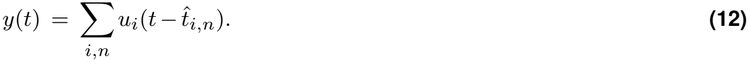

### 2.3 Phenomenological benchmark

The phenomenological model uses the same MUAP repertoire and summation (Eq. 12) but bypasses the spinal circuit: each MU fires at most once, and only the first *N*_*j*_ *≤ N*_MU_ MUs are recruited at the *j*-th TMS intensity. Their firing times are sampled from the rescaled cumulative distribution function (CDF) of a gamma distribution with shape *α* and rate *λ*, shared across intensities, with an additional time lag *t*_lag_ between intensities that mimics the input-dependent time to reach firing threshold, plus a common axonal delay *d*_axon_ (Supplementary Figure S1). Because the number of recruited MUs *N*_*j*_ at each TMS intensity is fitted as an independent free parameter rather than determined by spinal dynamics and DI-wave input, the phenomenological model is more flexible than the spinal-peripheral models and serves as their benchmark.

### 2.4 Model parameterization

The spinal-peripheral model comprises *N*_MU_ MNs and 2*N*_MU_ connection weights between the MNs and the RC population. To prevent overfitting, careful selection of free parameters is essential to balance model complexity and output flexibility. Free parameters were therefore selected iteratively by identifying sensitive parameters and their valid ranges. We considered two variants of the spinal-peripheral model: the full model with RC recurrent inhibition (RC^+^) and a reduced model without it (RC^*−*^). The RC^+^ model has 12 free parameters (Table 2). Parameters *p*_1*−*5_ determine the MN membrane resistances at five control points, the 1st, 10th, 20th, 60th, and 100th MNs, as cumulative sums of non-negative increments:

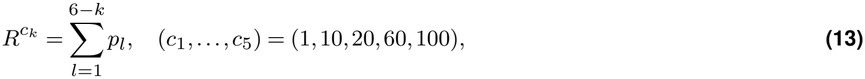

**Table 2.**
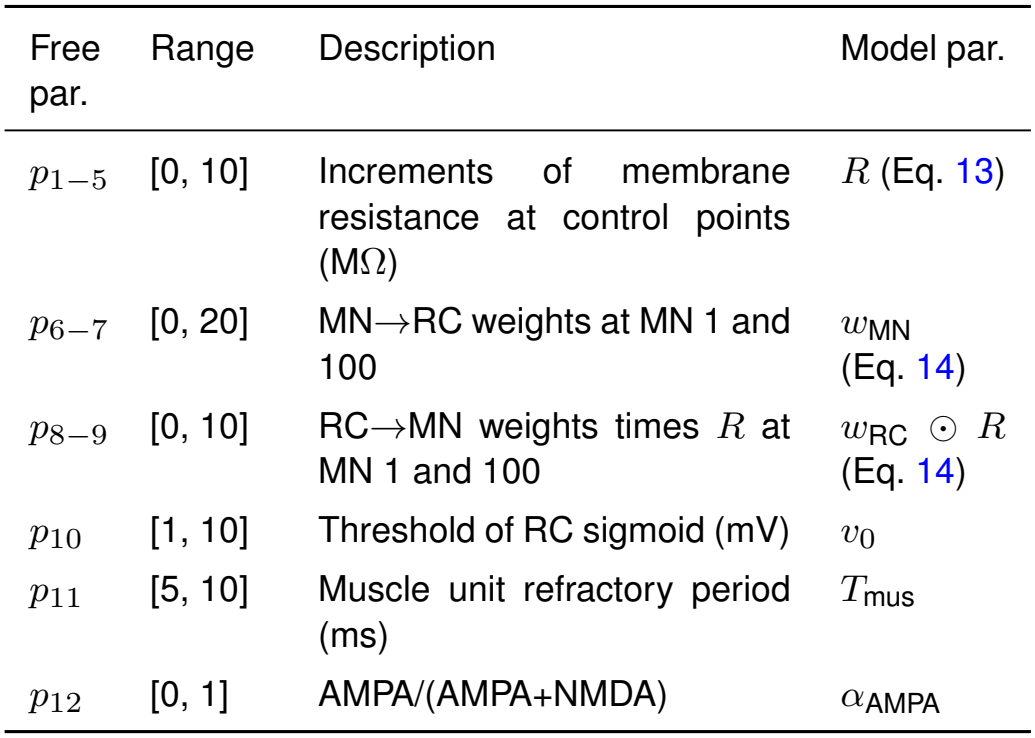
Free parameters of the spinal-peripheral RC^+^ model.

such that *R*^100^ = *p*_1_ and 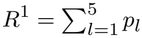. This construction guarantees that *R* decreases monotonically with the MN index, and the remaining values are interpolated using the Piecewise Cubic Hermite Interpolating Polynomial (PCHIP), which preserves monotonicity. Parameters *p*_6*−*9_ define linear profiles across the MN index for the MN*→*RC weights *w*_MN_ and for the RC*→*MN weights scaled by the membrane resistance, *w*_RC_ *⊙ R*, where *⊙* denotes element-wise multiplication:

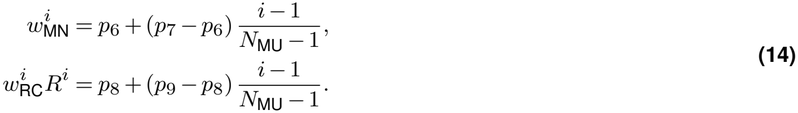

Because the RC current 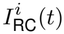 enters the membrane equation scaled by *R*^*i*^ (Eqs. 1 and 4), 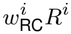 represents the effective strength of recurrent inhibition onto the *i*-th MN. This parameterization thus encompasses reports ranging from a uniform distribution of recurrent inhibition across MNs (*p*_8_ = *p*_9_) [35, 36] to inhibition that scales with MN size or recruitment order (*p*_8_ ≠ *p*_9_) [37–39]. The remaining free parameters are *v*_0_ (*p*_10_), *T*_mus_ (*p*_11_), and *α*_AMPA_ (*p*_12_). The RC^*−*^ model omits the RC population and its associated parameters (*p*_6*−*10_) and thus has 7 free parameters.

Simulated MEP amplitudes are rescaled to the maximum of the recorded MEPs, and *d*_axon_ *≥* 0 is set to the difference between simulated and recorded peak times at the strongest TMS intensity. The phenomenological model has 3 + *M* free parameters (Table 3): *α, λ, t*_lag_, and *N*_*j*_ for each of the *M* TMS intensities, with *d*_axon_ obtained in the same manner.

**Table 3.** Free parameters of the phenomenological model.

| Free par. | Range | Description | Model par. |
|---|---|---|---|
| *p*<sub>1</sub> | [0.1, 5] | Shape of gamma distribution | *α* |
| *p*<sub>2</sub> | [0.1, 5] | Rate of gamma distribution | *λ* |
| *p*<sub>3</sub> | [0, 2] | Time lag between TMS intensities (ms) | *t*<sub>lag</sub> |
| *p*<sub>4−∗</sub> | [4, *N*<sub>MU</sub>] | Number of recruited MUs at the *j*-th TMS intensity, *j* = 1, . . . , *M* | *N*<sub>*j*</sub> |

### 2.5 Model fitting and statistical analysis

Free parameters were optimized by minimizing the pointwise difference between simulated and recorded MEP waveforms, sampled at 0.1 ms intervals within 20–50 ms after the TMS pulse, across all TMS intensities. A genetic search algorithm was combined with the Gauss-Newton method, with parameters constrained to the ranges in Tables 2 and 3, following [40, 41]. GoF was quantified as *R*^2^ for MEP waveforms and for IO curves. Differences among the three models were assessed with Friedman tests (effect size: Kendall’s *W*), followed by post-hoc Wilcoxon signed-rank tests (effect size: rank-biserial correlation *r*_rb_) with Bonferroni correction (adjusted *α* levels: * < 0.0167, ** < 0.0033). Fitted parameters of the RC^+^ and RC^*−*^ models were compared using Wilcoxon signed-rank tests.

## 3 Results

### 3.1 Phenomenological benchmark

We first evaluated the phenomenological benchmark, in which MU firing times are sampled from the CDF of a gamma distribution (see Methods). The model assumes that MU firing times follow the same gamma distribution across TMS intensities, while the number of recruited MUs may differ. The phenomenological model captured both the MEP waveforms (average waveform *R*^2^ = 0.91) and the IO curves (average IO *R*^2^ = 0.96) across the ten participants (Supplementary Figure S2), demonstrating that the MUAP repertoire derived from the hand muscle model [33] is largely sufficient to account for individual variability in MEP waveforms. Owing to its simplifying assumptions, however, the benchmark has two properties that should be kept in mind when comparing it with the spinal-peripheral models. First, because only the first firing time of each recruited MU is modeled, the model emphasizes early MEP characteristics, such as peak latency and peak-to-peak amplitude, and is less precise for the overall MEP waveform. Late MEP components, such as the secondary peaks at higher TMS intensities in participant S1, which may arise from repeated MU firing, are therefore not well reproduced. Second, because *N*_*j*_ is fitted as an independent free parameter at each TMS intensity, the model is not constrained to a monotonic IO relationship and can overfit irregular IO responses, as in participant S8. Its IO *R*^2^ may therefore overestimate what a physiologically constrained model can achieve. Thus, the benchmark indicates how well the MUAP repertoire alone can reproduce the recorded MEPs, whereas the spinal-peripheral models must generate MU firing patterns from the DI-wave input through spinal dynamics.

### 3.2 RC inhibition shapes MEP waveforms by regulating MN firing

As demonstrated by the phenomenological benchmark (Section 3.1), MU firing times can largely account for inter-individual variability in MEP waveforms, because the MUAP repertoire is fixed. In the spinal-peripheral model, MU firing times are in turn determined by MN spike timing, which depends on participant-specific parameters such as the MN membrane resistance (*R*) and the MN-RC connectivity (*w*_MN_ and *w*_RC_, Table 2). Note that inter-individual variability in MEP waveforms was attributed largely to participant-specific model parameters rather than to the DI-wave input, which was reconstructed from the same group-level DI-wave IO curves for all participants and differed only in the individual RMT used to scale TMS intensities (Figure 2).

We examined the waveform fits from two perspectives: the overall waveform shape, including late components such as secondary peaks, and the nearly constant MEP peak latency across TMS intensities. Figure 4 illustrates both for a representative participant (S1). Regarding the overall shape, the phenomenological model (Figure 4A, *R*^2^ = 0.87), in which each MU fires at most once, reproduced the peak amplitudes and latencies but missed the secondary peaks at 50%, 53%, and 56% MSO. Such late components require repeated MU firing distributed across the MEP period, which both spinal-peripheral models can generate. However, the RC^+^ model (Figure 4B, *R*^2^ = 0.92) reproduced these peaks more closely than the RC^*−*^ model (Figure 4C, *R*^2^ = 0.87), indicating that repeated MU firing alone is not sufficient and that RC inhibition shapes its temporal distribution.

**Figure 4.**
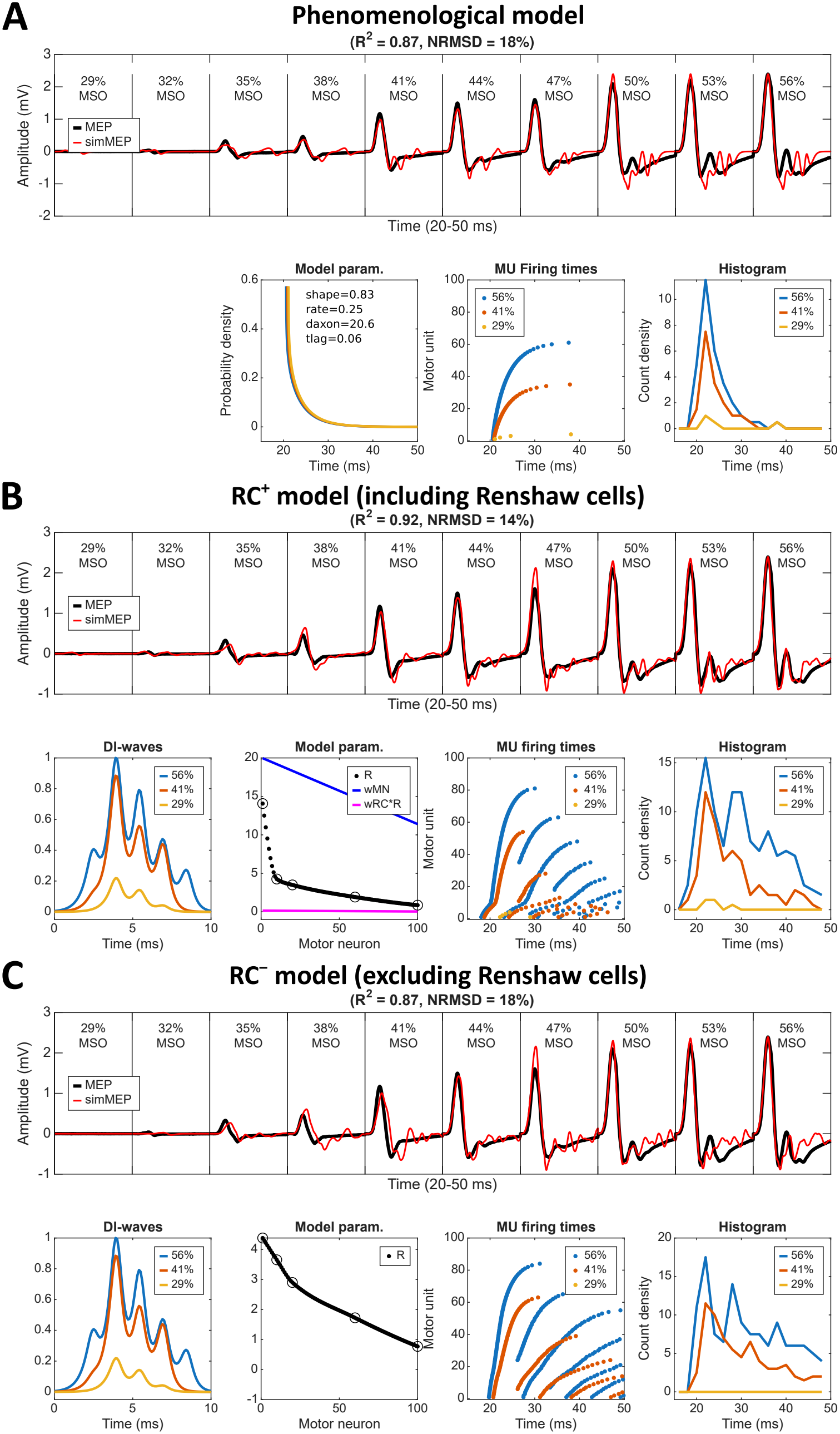
Fitting results of a representative participant (S1) using three models. The three models, including **(A)** phenomenological model, **(B)** spinal-peripheral RC^+^ model, and **(C)** spinal-peripheral RC^*−*^ model, are fitted to the MEP waveforms of participant S1 under various TMS intensities from 29% to 56% MSO. The first row shows the MEP waveforms (from 20 to 50 ms after the TMS pulse) for a visual comparison between the recordings (black trace) and simulations (red trace). The second row, from left to right, displays the model input (DI-waves to the spinal-peripheral models), the fitted model parameters, the MU firing times, and the histogram of MU firing times under three example TMS conditions (29%, 41%, and 56% MSO).

Regarding peak latency, the phenomenological model reproduced the nearly constant peak latency only through physiologically implausible, near-zero fitted time lags (*t*_lag_ = 0.06 ms for S1; approaching zero in S2, S5, S6, and S9, Supplementary Figure S2), whereas firing latency should vary with input strength. In the spinal-peripheral models, weaker TMS produces DI-waves with lower amplitudes and later onsets (e.g., 29% vs. 56% MSO), which would predict later MU firing. In the RC^+^ model, however, the first peak of the MU firing-time histogram remains aligned across intensities: MU firing begins later under weaker stimulation but also ceases earlier, yielding a stable MEP peak latency (Figure 4B). In the RC^*−*^ model, without this inhibitory feedback, the first peak fails to decay sufficiently early (e.g., at 41% MSO), producing delayed MEP waveforms at lower TMS intensities and a poorer overall match (Figure 4C). Together, these results indicate that RC inhibition contributes to both the overall shape and the stable peak latency of MEP waveforms.

To understand how RC inhibition regulates MN firing, we examined the internal dynamics underlying a single MEP waveform of participant S1 at 56% MSO (Figure 5). In the RC^+^ model (Figure 5A), the effective excitatory conductance of MN1 follows the DI-wave input, whereas the effective inhibitory conductance rises as the RC population becomes active within approximately 10 ms after TMS onset, progressively counterbalancing the excitatory drive. The RC^*−*^ model (Figure 5B) compensates for the absence of inhibitory feedback by a higher AMPA contribution (*α*_AMPA_ = 0.55), which shortens the excitatory drive and yields a superficially similar pattern of MU firing times. Across participants, the RC^*−*^ model compensated through an elevated *α*_AMPA_ (all participants) and lower resistances *R* of small MNs (Section 3.3.2). However, *α*_AMPA_ and *R* alone provide insufficient flexibility to reproduce the full range of MEP waveform shapes across TMS intensities.

**Figure 5.**
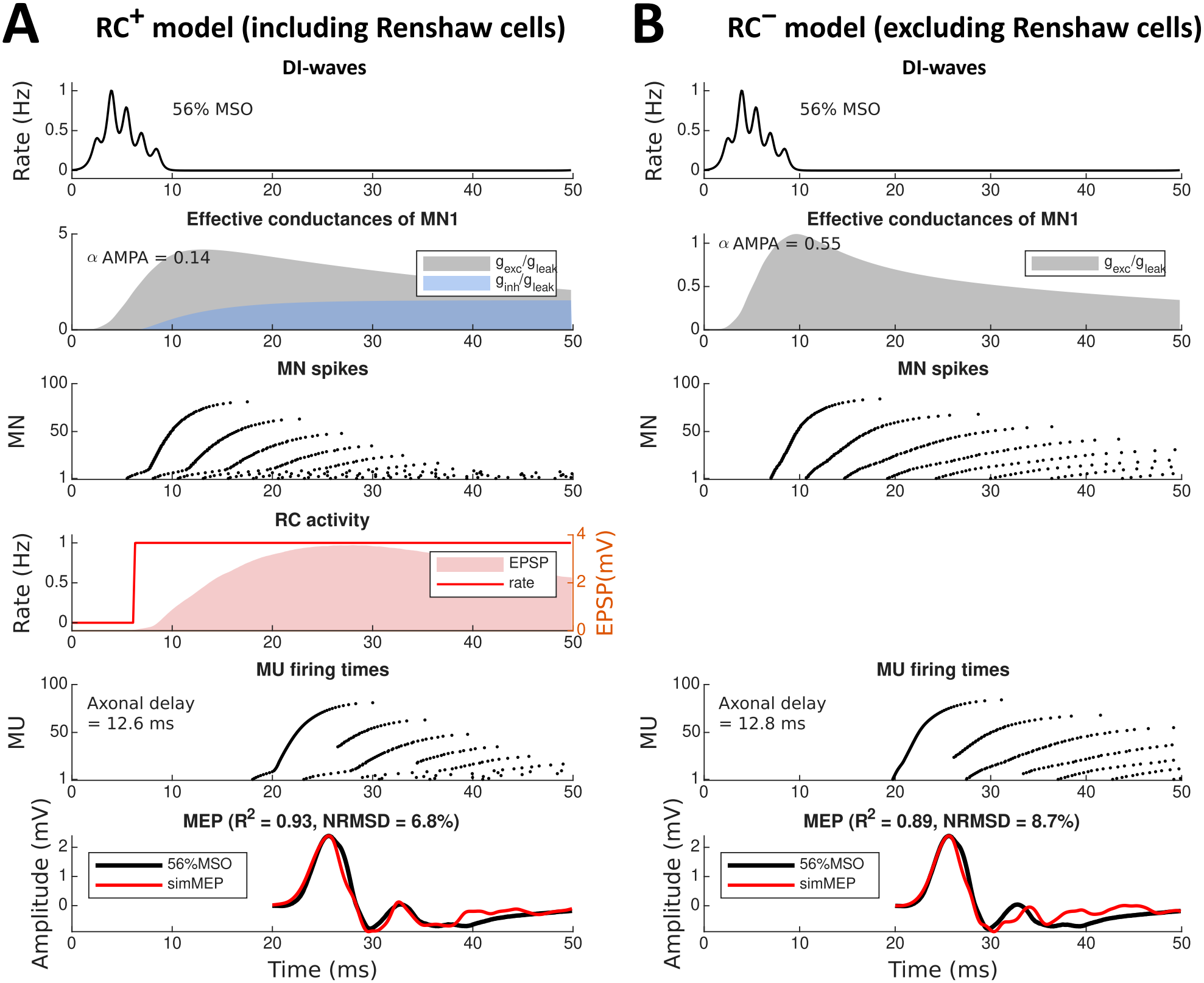
Detailed process of MEP generation. A single MEP waveform of participant S1 at 56% MSO (corresponding to Figure 4B and C) is simulated using **(A)** the spinal-peripheral RC^+^ model and **(B)** the spinal-peripheral RC^*−*^ model. The deconvolved DI-waves (black trace, 1st row) drive the MNs, increasing the effective excitatory conductance *g*_exc_*/g*_leak_ = *R*^1^ · (*g*_DI_ ⊗ *m*_DI_) (*t*) (gray shade, 2nd row; shown for MN1 only, with the fitted *α*AMPA indicated) and generating MN spikes (black dots, 3rd row). In the RC^+^ model **(A)**, the MN spikes increase the EPSP (*v*_RC_) of the RC population (red shade, right axis, 4th row) and activate it (red curve, firing rate, left axis), which in turn increases the effective inhibitory conductance *g*_inh_*/g*_leak_ = *R*^1^ *·* (*g*_RC_ *⊗ m*_RC_) (*t*) of the MNs (blue shade, 2nd row), with *R*^1^ = 1*/g*_leak_ (Eqs. 1–4). In both models, the MN spikes reach the muscle units after the axonal delay and generate MUAPs whenever they arrive outside the muscle unit refractory period, yielding the MU firing times (black dots, 5th row). The simulated MEP (simMEP, red trace, 6th row) is the sum of the time-shifted MUAPs and is compared with the recorded MEP (black trace). *R*^2^ and the normalized root-mean-square deviation (NRMSD, the root-mean-square difference between the recorded and simulated waveforms divided by the peak-to-peak range of the recorded waveform) quantify the GoF for this single waveform.

### 3.3 Group-level model comparison

We next compared the three models across all participants. In addition to testing whether the RC^+^ model performs as well as the phenomenological benchmark, we tested whether the simpler RC^*−*^ model could perform as well as the RC^+^ model, as recurrent inhibition might play a limited role in fast, transient events such as MEPs. This comparison is valuable because the RC^*−*^ model has a simpler structure, which potentially reduces model complexity and mitigates the risk of overfitting. The GoF of the three models is summarized in Figure 6, and the individual fits of the RC^+^ and RC^*−*^ models are shown in Supplementary Figures S3 and S4, respectively.

**Figure 6.**
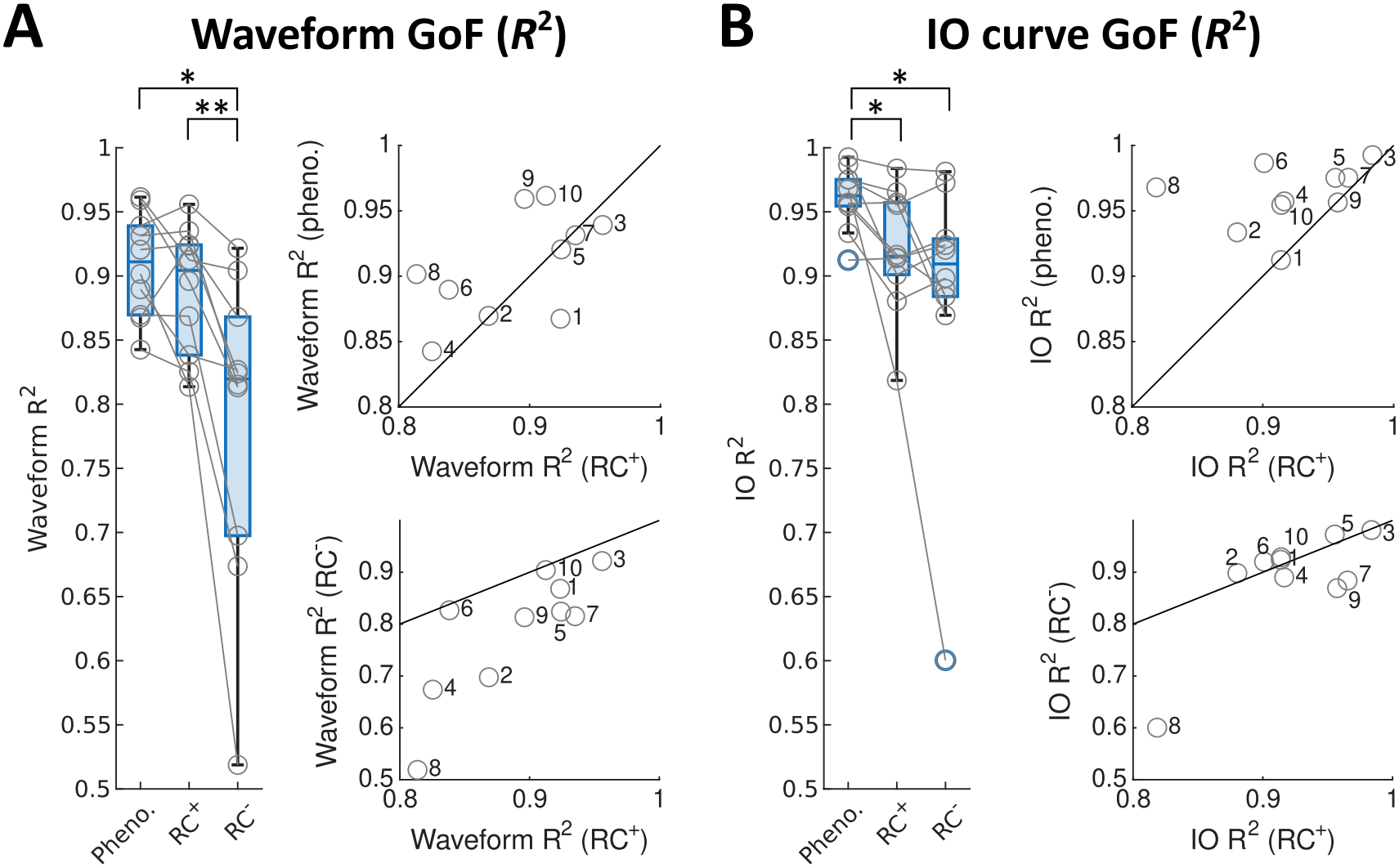
Waveform and IO curve GoF across the three models. **(A)** GoF (*R*^2^) based on MEP waveforms. **(B)** GoF (*R*^2^) based on MEP IO curves. The three models include the phenomenological model (Pheno.), the spinal-peripheral RC^+^ model, and the spinal-peripheral RC^*−*^ model. The box plots indicate the median, lower and upper quartiles, and the minimum and maximum values. Gray circles represent individual participants. Numbers next to gray circles denote participant IDs. In each panel, the upper scatter plot compares the RC^+^ model (x-axis) with the phenomenological model (y-axis), and the lower scatter plot compares the RC^+^ model (x-axis) with the RC^*−*^ model (y-axis). Asterisks denote significant post-hoc Wilcoxon signed-rank tests (Bonferroni-corrected *α* levels: * < 0.0167, ** < 0.0033).

#### 3.3.1 Waveform and IO goodness-of-fit

Compared with the phenomenological model (average waveform *R*^2^ = 0.91), the RC^+^ model achieved comparable performance (0.89), whereas the RC^*−*^ model performed worse (0.79). The three models differed significantly in waveform GoF (Figure 6A, Friedman test: *χ*^2^(2) = 12.20, *p* = 0.0022, Kendall’s *W* = 0.61, large effect). Post-hoc tests revealed no significant difference between the phenomenological and RC^+^ models (*p* = 0.3223, *r*_rb_ = 0.38), whereas the RC^*−*^ model performed significantly worse than both the phenomenological model (*p* = 0.0039, *r*_rb_ = 0.96) and the RC^+^ model (*p* = 0.0020, *r*_rb_ = 1.00). The waveform *R*^2^ of the RC^+^ model exceeded that of the RC^*−*^ model in all 10 participants (Figure 6A, lower scatter plot; individual fits in Supplementary Figures S3 and S4). With the RC^+^ model, five participants achieved waveform *R*^2^ values above 0.9 and the remaining five between 0.8 and 0.9, with the lowest value in S8 (*R*^2^ = 0.81), likely due to irregular IO responses. Moreover, the GoF was positively correlated across the three models (Figure 6, scatter plots), suggesting that the GoF partly depends on how individual hand muscles deviate from the MUAP repertoire, with exceptions such as S1 (complex waveforms poorly represented by the benchmark) and S8 (benchmark overfitting). For IO GoF (Figure 6B), the main effect of model type was also significant (*χ*^2^(2) = 7.40, *p* = 0.0247, *W* = 0.37, medium effect): the phenomenological model (average IO *R*^2^ = 0.96) outperformed both the RC^+^ model (0.92; *p* = 0.0098, *r*_rb_ = 0.89) and the RC^*−*^ model (0.89; *p* = 0.0098, *r*_rb_ = 0.89), while the two spinal-peripheral models did not differ (*p* = 0.4922, *r*_rb_ = 0.27). The IO GoF can be misleading because it reflects only peak-to-peak amplitudes: a minor time shift decreases waveform accuracy while leaving the amplitude unchanged. Accordingly, although the RC^*−*^ model yielded lower waveform *R*^2^ in every participant, its IO *R*^2^ was slightly higher in S1, S2, S5, S6, and S10. In S2, for example, the RC^*−*^ model produced distorted MEP waveforms (*R*^2^ = 0.7) despite a high IO fit (*R*^2^ = 0.9) (Supplementary Figures S3 vs. S4).

In summary, the RC^+^ model matched the benchmark in waveform GoF, whereas the RC^*−*^ model did not. Moreover, the models were more strongly differentiated by waveform GoF (*W* = 0.61) than by IO GoF (*W* = 0.37), indicating that waveform analysis carries more discriminative information than amplitude-based measures alone.

#### 3.3.2 Fitted parameters

In the RC^+^ model (Supplementary Figure S3, column 3), the fitted membrane resistances declined sharply near MN20 in all participants, marking the transition from small to large MNs. The MN-RC connectivity (*w*_MN_ and *w*_RC_) varied substantially across participants, with no consistent positive or negative trend.

In the RC^*−*^ model, the absence of RC inhibition was compensated by other parameters (Supplementary Figure S4, column 3): *α*_AMPA_ was higher than in the RC^+^ model in all participants (RC^+^ vs. RC^*−*^: median 0.31 vs. 0.91, *p* = 0.0020), and the mean membrane resistance of small MNs (MN1–20) was lower (median 11.69 vs. 6.25 MΩ, *p* = 0.0195). In contrast, the axonal delay (median 15.25 vs. 14.20 ms, *p* = 0.2383) and the muscle unit refractory period (median 5.02 vs. 5.32 ms, *p* = 0.3750) did not differ between the two models. Thus, the RC^*−*^ model compensated mainly through synaptic and MN properties (*α*_AMPA_ and *R*), whereas the axonal and muscle unit properties remained unchanged.

#### 3.3.3 MU firing times

In the RC^+^ model, the estimated MU firing times (Figure 4B; Supplementary Figure S3, column 4) show that stronger TMS recruits more MUs, which fire earlier and repetitively for longer. The recruitment of additional, higher-threshold MUs results from the temporal summation of successive descending volleys (Figure 5). In the model, this summation occurs at the level of synaptic conductance: each volley is filtered by the AMPA and NMDA kernels (Eq. 5), and the slow decay of the NMDA component allows the effects of successive volleys to accumulate, while RC inhibition limits the resulting discharge. Through these mechanisms, the high-frequency (approximately 670 Hz) corticospinal input is transformed into much lower-frequency MU discharges. The duration of MU firing varied across participants (prolonged in S1, S2, S3, S6, and S9; brief in S4 and S8) and was not always directly reflected in the MEP waveform, suggesting that the MU firing times and fitted parameters may provide valuable information for characterizing individual profiles.

In the RC^*−*^ model, the parameter compensations altered the MU firing patterns (Supplementary Figure S4, column 4): MU firing persisted longer than in the RC^+^ model (S1, S2, S7, S8, and S9) or, owing to low *R*, was insufficient to produce MEPs at weak intensities (S4 and S5), yielding substantially lower waveform *R*^2^. Overall, the differences in GoF, fitted parameters, and MU firing times indicate that the spinal-peripheral model incorporating RC inhibition is the more appropriate choice for modeling MEPs.

## 4 Discussion

We developed a spinal-peripheral model of TMS-evoked MEPs and evaluated it on MEP waveforms from 10 healthy participants. The RC^+^ model performed as well as a flexible phenomenological benchmark in waveform GoF, reproducing individual MEP waveforms over a wide range of TMS intensities with an average GoF of approximately 0.9, whereas the RC^*−*^ model showed significantly lower waveform GoF. For IO curves, the benchmark outperformed both spinal-peripheral models, as expected given its independently fitted *N*_*j*_ at each TMS intensity. The RC^+^ model outperformed the RC^*−*^ model in waveform GoF in every participant (rank-biserial *r*_rb_ = 1.00). These results underscore the critical role of RC recurrent inhibition in regulating MN spike timing, which in turn shapes the temporal features of the MEP. Combining LIF MNs and a rate-based RC population with a physiologically grounded MUAP repertoire, the model is both computationally efficient and biologically plausible, making it a promising candidate for integration with motor cortex models that generate DI-waves, toward individualized models of the TMS-evoked motor pathway.

### 4.1 Latent activity and fitted parameters

Fitting the model to individual MEP waveforms enables inference of latent neural dynamics that are not directly observable, including MN spike trains, RC activity, and MU firing times, together with interpretable parameters that capture participant-specific properties of the corticospinal pathway, including MN membrane resistance, MN-RC connectivity, synaptic AMPA/NMDA balance, axonal conduction delay, and muscle unit refractoriness. Reproducing late MEP components (>30 ms after stimulation) required sustained MN firing beyond the initial discharge, which the single-activation phenomenological model cannot provide (Figure 4), despite its higher waveform *R*^2^ in six participants (Figure 6A) owing to its independently adjustable *N*_*j*_ (Table 3). These results suggest that MEP waveforms acquired across a wider range of TMS intensities can provide useful constraints for model fitting. In the RC^+^ model, sustained firing was supported by a larger NMDA contribution (lower *α*_AMPA_), and the fitted muscle unit refractory periods *T*_mus_ tended toward small values (Section 3.3.2), allowing repeated MUAPs within a single MEP, consistent with experimental observations of repeated MU firing [42]. During parameter selection, we observed that a steep slope *r* of the RC sigmoid was necessary for recurrent inhibition to act in time: because the glycinergic RC*→*MN kernel already introduces a delay (*τ*_*r*_ = 1 ms, *τ*_*d*_ = 6 ms), a shallow sigmoid would further delay the inhibition and fail to suppress MN firing promptly. We therefore fixed *r* at 10 mV^*−*1^, yielding a nearly step-like RC response (Figure 5), in line with the observation that RCs respond fast and can be activated by a single MN spike [32].

Including RC inhibition yielded a broader range of MN membrane resistances, with higher resistances among the smallest MNs (MN1–20) and thus a steeper decline of resistance with MN index (Section 3.3.2; Supplementary Figure S5). Combined with recurrent inhibition, this broader range enables more diverse MN recruitment patterns across TMS intensities. Given the inverse relationship between membrane resistance and cell size, this distribution provides an indirect estimate of the MN size spectrum, a property linked to motor function and aging [43–46]. As input and output amplitudes are normalized during fitting, these estimates should be interpreted in relative terms. The estimated axonal delays (median approximately 15 ms) fall within biologically plausible ranges [47, 48], as do the muscle unit refractory periods (approximately 5 ms), and neither differed between the two models (Section 3.3.2). In contrast, the substantially lower *α*_AMPA_ in the RC^+^ than in the RC^*−*^ model reflects distinct strategies: in the RC^+^ model, slow NMDA currents provide sustained excitatory drive that RC inhibition regulates, whereas the RC^*−*^ model limits the duration of MN excitation through a higher AMPA weight, consistent with the different contributions of non-NMDA and NMDA receptors to EPSP amplitude and duration in motoneurons [49]. Note that *α*_AMPA_, a mixing weight between peak-normalized kernels (Eq. 5), is not directly comparable to experimentally reported AMPA/NMDA ratios, as the model does not include the voltage-dependent Mg^2+^ block of NMDA receptors. Moreover, because the AMPA and NMDA kernels are normalized to equal peak conductance (Eq. 7), the effective charge ratio between the two components also depends on their time constants, which should be accounted for when relating *α*_AMPA_ to physiological AMPA/NMDA ratios.

Not all parameters are equally constrained by the data. The MN-RC connectivity parameters (*p*_6*−*9_ in Table 2) varied substantially across participants, reflecting a degree of non-identifiability: MN spike timing is jointly determined by *R, w*_MN_, *w*_RC_, *v*_0_, and *α*_AMPA_, leading to non-unique solutions. Additional physiological constraints on MN-RC connectivity may help reduce this degeneracy. Clinically, MEPs are typically summarized by a few scalar measures, such as MEP presence, motor threshold, and amplitude, which are interpreted as markers of corticospinal integrity or cortical excitability. For example, MEP presence early after stroke predicts upper limb recovery [3, 4], whereas reduced motor thresholds and increased MEP/CMAP amplitude ratio early in the disease indicate cortical hyperexcitability in amyotrophic lateral sclerosis [6, 50, 51]. Because such measures reduce the MEP waveform to its amplitude or presence, alterations at different stages of the motor pathway may yield similar values. Fitting the model to patient MEP waveforms could complement these measures by estimating spinal and peripheral properties, such as MN excitability, recurrent inhibition, and axonal conduction delay, as well as the underlying MU firing patterns that shape the MEP waveform. Such estimates may help attribute clinical MEP abnormalities to specific stages of the motor pathway.

### 4.2 Model complexity

We aimed to simplify the model architecture while preserving sufficient flexibility to reproduce individual MEP waveforms and IO characteristics. Our model lies between two classes of existing models. Previous MEP models [12, 13] restricted the spinal component to MN populations and represented surface EMG as a superposition of MUAPs parameterized by Hermite-Rodriguez functions [52, 53]. They targeted cortical phenomena reflected mainly in MEP amplitude, such as short- and long-interval intracortical inhibition, intracortical facilitation, and the cortical silent period. Because fine temporal waveform structure was not their primary objective, RC inhibition was not incorporated. Our results show that such a spinal component is insufficient to reproduce the temporal features of MEP waveforms, motivating the inclusion of an RC population in our model. Conversely, models focused on RC function have employed large spinal networks, such as 300 *α*MNs and 300 RCs [18, 54] or 377 MNs and 64 RCs [19, 55], supported by evidence that a single MN input reliably drives RC firing [32]. These models target sustained behaviors such as force production, tremor, and corticomuscular coherence. For the brief, transient MEP evoked by a single TMS pulse, a single lumped RC population was sufficient. Together, these comparisons suggest that the complexity of our model is appropriate for the task: RC inhibition is required to reproduce the fine temporal structure of MEP waveforms, whereas a single lumped RC population avoids the computational overhead and risk of overfitting associated with larger-scale spinal configurations, balancing biological plausibility with the computational tractability needed for individual-level fitting.

### 4.3 Limitations

First, the DI-wave input was reconstructed from group-level DI-wave IO curves measured with PA coil orientation (as in the MEP recordings included in this study). Other orientations, such as anterior-posterior and latero-medial, alter the induced electric field [56] and the underlying cortical circuit dynamics [1, 57], producing different DI-waves and MEPs [9]. Evaluating the model under multiple orientations would provide a more stringent test of its flexibility, and frameworks linking orientation-specific electric fields to cortical dynamics and DI-wave generation [56, 58, 59] could serve as upstream input for orientation-aware MEP simulation. Second, individualized DI-wave recordings, either invasively from the cervical epidural space or non-invasively from surface electrodes over the cervical spine, could improve the estimation of participant-specific parameters, although noisy individual DI-waves may also mislead parameter estimation. Future work will assess whether the estimated parameters reliably reflect individual structural properties at realistic noise levels and whether they are identifiable and useful for clinical applications.

## Data and Code Availability

The MEP dataset analyzed in this study is publicly available on the Open Science Framework (OSF) at https://osf.io/5ry92/, originally collected and described by Sorkhabi et al. [21]. The MATLAB and Python implementations of the models are available at https://github.com/vscChien/MEPmodeling and https://github.com/EleBern/MEPmodeling-Python, respectively.

## Author Contributions

V.C.: Conceptualization, Methodology, Software, Formal Analysis, Writing – Original Draft, Funding Acquisition. E.B.: Software, Validation, Writing – Review & Editing. E.M.: Writing – Review & Editing. P.W.: Software. M.L.: Software, Methodology, Formal Analysis, Writing – Review & Editing. J.L.: Software, Methodology, Formal Analysis, Writing – Review & Editing. Ka.W.: Investigation, Writing – Review & Editing. J.O’S.: Funding Acquisition, Supervision, Writing – Review & Editing. T.D.: Funding Acquisition, Supervision, Writing – Review & Editing. F.R.: Writing – Review & Editing. V.D.L.: Writing – Review & Editing. J.H.: Supervision, Writing – Review & Editing, Funding Acquisition. T.R.K.: Conceptualization, Supervision, Writing – Review & Editing, Funding Acquisition. Ko.W.: Conceptualization, Methodology, Supervision, Writing – Review & Editing, Funding Acquisition. H.S.: Conceptualization, Methodology, Supervision, Writing – Review & Editing, Funding Acquisition.

## Declaration of generative AI and AI-assisted technologies in the manuscript preparation process

During the preparation of this work the authors used Claude (Anthropic) in order to improve the wording and readability of the manuscript text. After using this tool, the authors reviewed and edited the content as needed and take full responsibility for the content of the published article.

## Supplementary Methods

### Extracellular potential kernel for DI-wave deconvolution

To generate the extracellular potential (EP) kernel used to deconvolve the reconstructed DI-waves (main text, Figure 2B), action potential profiles *V* (*t*) were simulated with an existing toolbox for myelinated axons [1]. The EP was computed from these profiles according to the following convolution integral [2]:

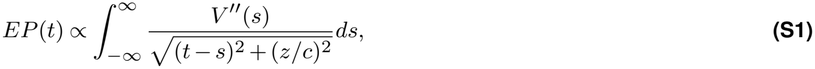

with a recording distance of *z* = 5 mm and a propagation velocity of *c* = 50 m/s.

### Phenomenological model

The phenomenological model generates MEP waveforms using the same MUAP repertoire {*u* (*t*)} and summation formula as the spinal-peripheral model, except that each MU fires at most once, with its firing time 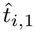 for *i* = 1, …, *N*_*j*_. Here, *N*_*j*_ *≤ N*_MU_ denotes the number of recruited MUs at the *j*-th TMS intensity, and MUs with index *i > N*_*j*_ are not recruited. The firing times 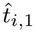 are derived from a gamma distribution characterized by a shape parameter *α* and a rate parameter *λ* (Figure S1A):

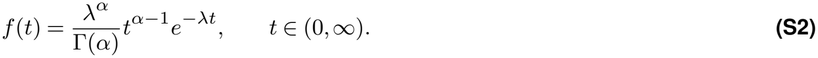

For a given TMS intensity condition *j*, the first firing times are allocated to *N*_*j*_ recruited MUs by sampling *N*_*j*_ values from the rescaled cumulative distribution function (CDF) of the gamma distribution (Figure S1B). The sorted firing times for different TMS intensities are derived by varying *N*_*j*_ while using the same CDF. To account for differences in the time required for a leaky integrate-and-fire (LIF) neuron to reach its firing threshold under varying synaptic input levels, an additional time lag *t*_lag_ is introduced between TMS conditions. A common axonal delay *d*_axon_ is then added to obtain the MU firing times.

Assuming a gamma distribution allows the phenomenological model to capture individual variability, accommodating patterns such as exponentially increasing and sigmoid-shaped firing times (Figure S1B).

**Figure S1.**
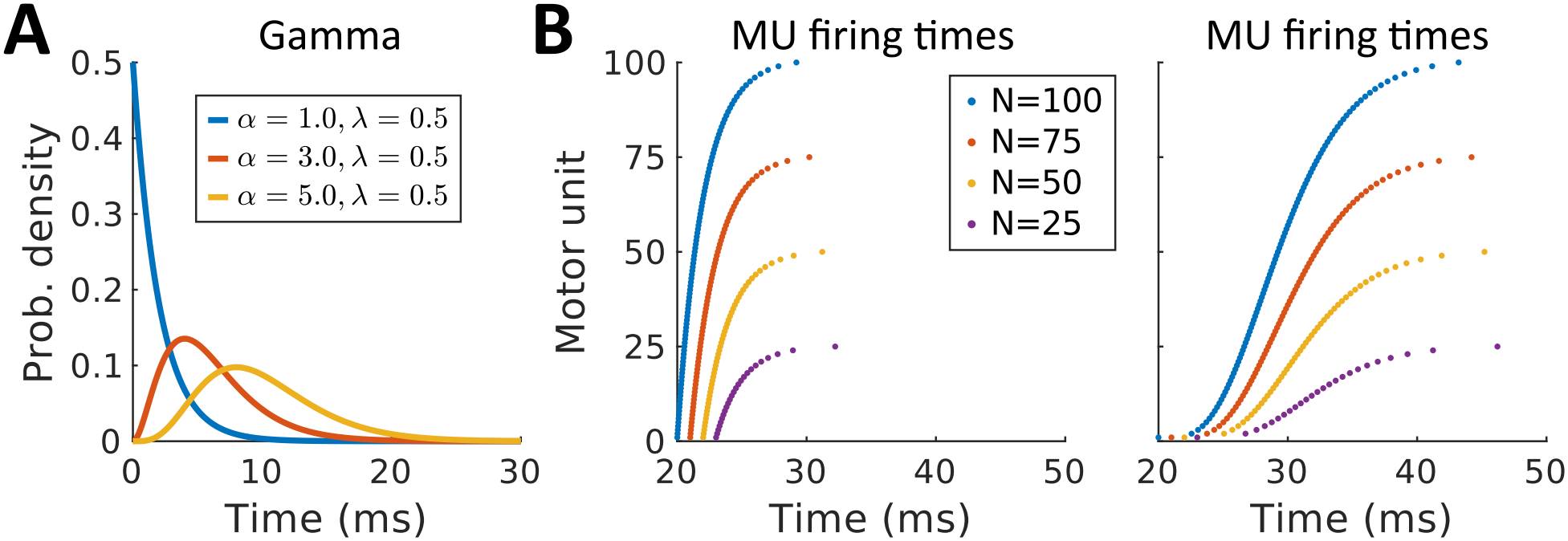
MU firing times derived from gamma distributions. **(A)** Gamma distributions defined by shape *α* and rate *λ* parameters. Two examples of MU firing times derived from the CDF of gamma distributions. The CDF is rescaled to the number of MUs (*N*_*j*_) recruited at TMS intensity *j*. Examples show *N*_*j*_ = 25, 50, 75, and 100, corresponding to MN recruitment from low to high TMS intensities. An additional time lag (*t*_lag_, e.g., 1 ms) between the TMS conditions accounts for differences in the time required for a leaky integrate-and-fire (LIF) neuron to reach firing threshold under different input levels. A common axonal delay (*d*_axon_, e.g., 20 ms) is then added on top of these firing times. The left example illustrates exponentially growing firing times ([*α, λ*] = [1, 0.5]), while the right example depicts sigmoid-shaped firing times ([*α, λ*] =[5, 0.5]). During fitting, free parameters (*α, λ, t*_lag_, and *N*_*j*_) are optimized to best match real MEP data, with *d*_axon_ calculated as the time difference between real and simulated MEP peaks.

## Supplementary Figures

**Figure S2.**
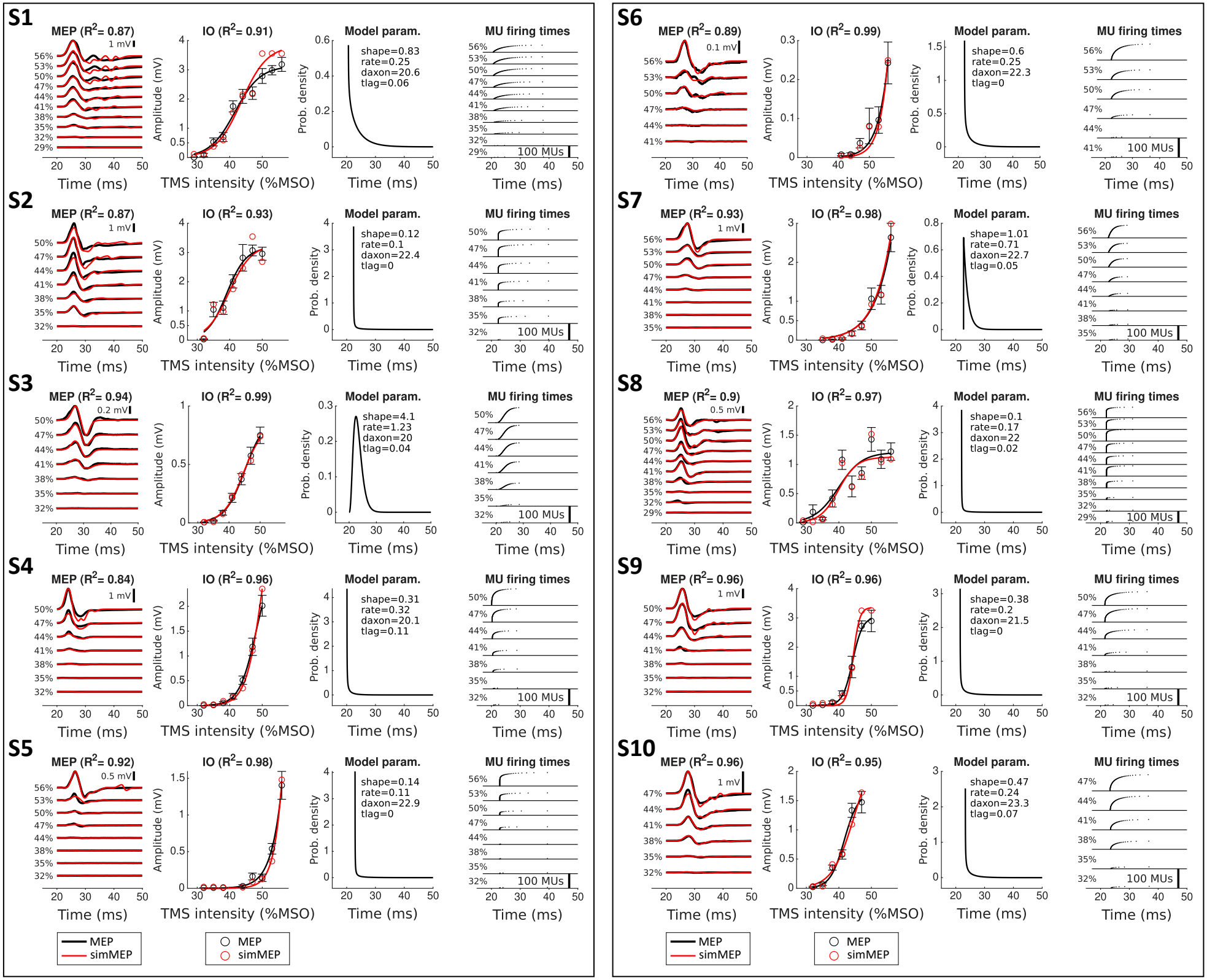
Phenomenological model fitted to the MEPs of individual participants. Column 1: Average motor-evoked potentials (MEPs) from 15 trials at different transcranial magnetic stimulation (TMS) intensities (%MSO, black traces) compared with MEPs simulated by the phenomenological model (red traces). Goodness-of-fit (GoF) of MEP waveforms is quantified by *R*^2^ value. **Column 2**: MEP input-output (IO) responses showing the peak-to-peak amplitudes of experimental MEPs (black circles, mean*±*SE, *n* = 15) and simulated MEPs (red circles) plotted against TMS intensity. The *R*^2^ value indicates the GoF of the IO curve. IO curves (black and red) are fitted sigmoid functions for visual comparison. **Column 3**: Fitted model parameters, including the gamma distribution of MU firing times (black curve) defined by the shape (*α*) and rate (*λ*) parameters, shifted by the axonal delay (*d*_axon_). **Column 4**: Estimated first firing times of 100 MUs across TMS intensities.

**Figure S3.**
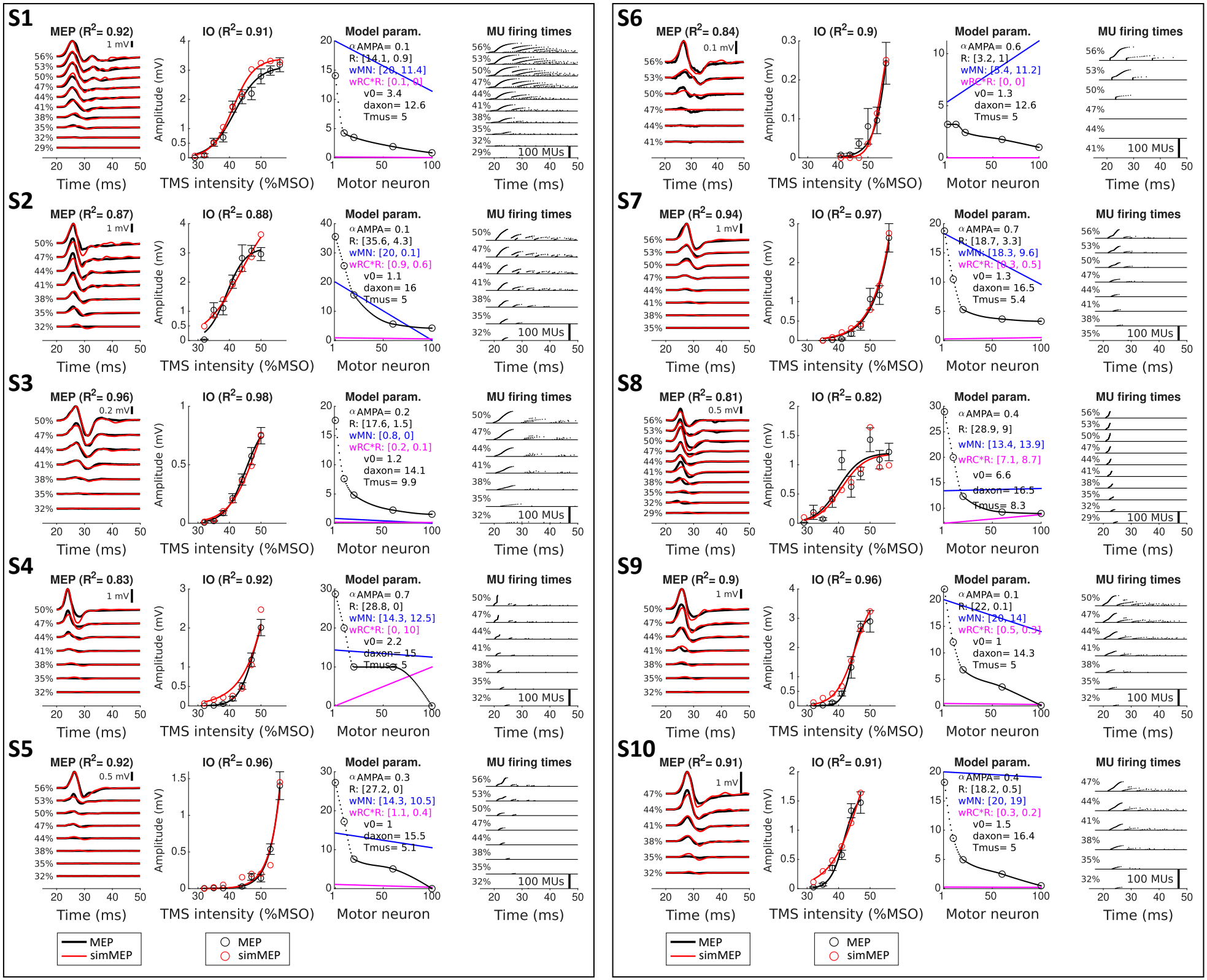
Spinal-peripheral RC^+^ model fitted to the MEPs of individual participants. Column 1: Average motor-evoked potentials (MEPs) from 15 trials at different transcranial magnetic stimulation (TMS) intensities (%MSO, black traces) compared with MEPs simulated by the spinal-peripheral RC^+^ model (red traces). Goodness-of-fit (GoF) of MEP waveforms is quantified by *R*^2^ value. **Column 2**: MEP IO responses showing the peak-to-peak amplitudes of experimental MEPs (black circles, mean*±*SE, *n* = 15) and simulated MEPs (red circles) plotted against TMS intensity. The *R*^2^ value reflects the GoF between experimental and simulated data. Sigmoid curves (black and red) are fitted for visual comparison. **Column 3**: Fitted model parameters, including membrane resistances of 100 MNs (black dots smoothly interpolated from five circles: MNs 1, 10, 20, 60, and 100) and linear connection strengths from MNs to the RC population (*w*_MN_, blue) and from the RC population back to MNs multiplied by the membrane resistance (*w*_RC_ *⊙ R*, magenta). The values in the brackets indicate these values for the 1^st^ and 100^th^ MNs, respectively. **Column 4**: Estimated MU firing times across TMS intensities.

**Figure S4.**
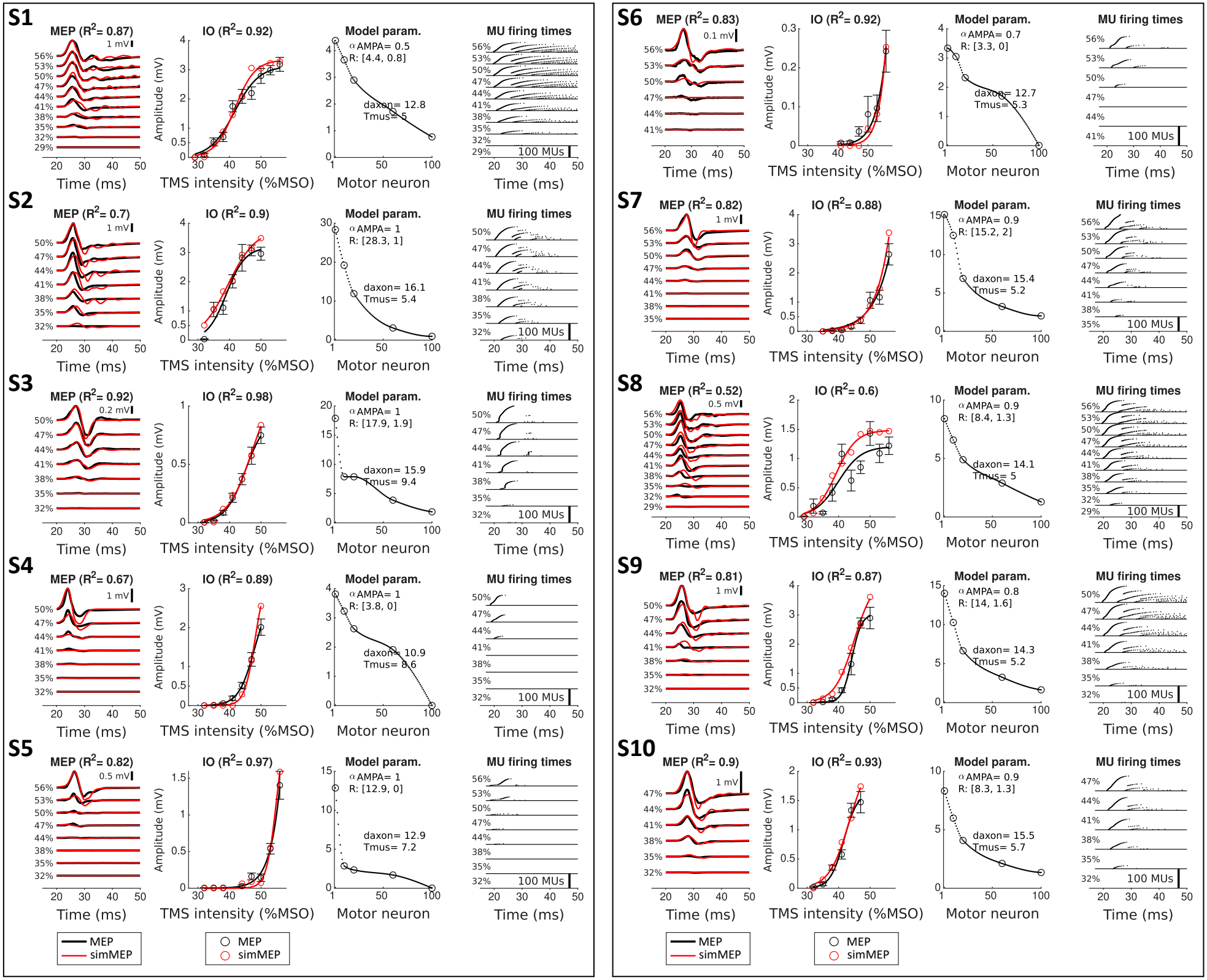
Spinal-peripheral RC^*−*^ model fitted to the MEPs of individual participants. The same arrangement as in Supplementary Figure S3, with the exception that in column 3, the parameters do not include connections with the RC population and RC firing threshold.

**Figure S5.**
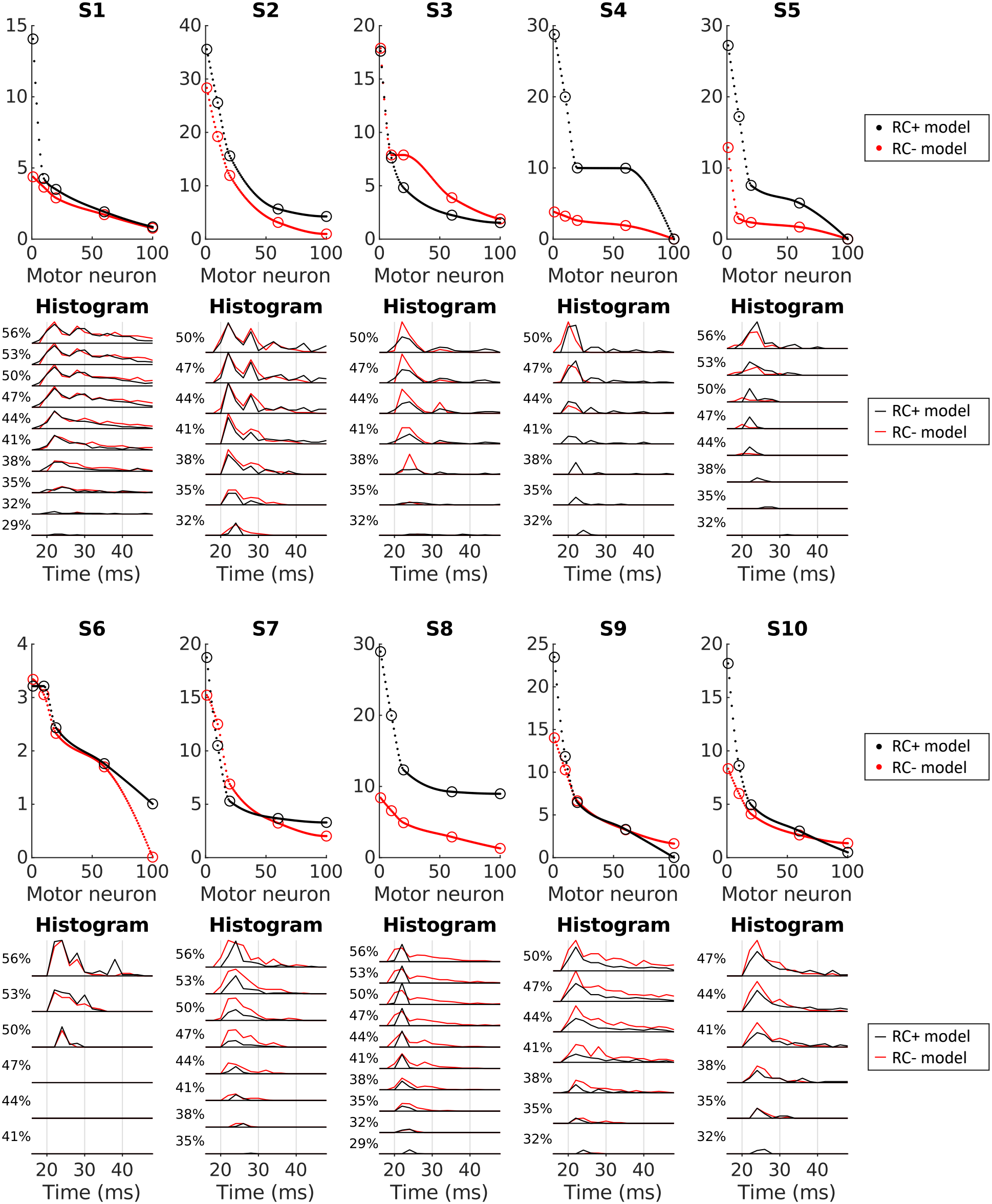
Fitted membrane resistances and motor unit (MU) firing times in the spinal-peripheral RC^+^ and RC^*−*^ models. (1st and 3rd rows) Fitted membrane resistances *R* (MΩ) of the 100 MNs in the RC+ model (black) and the RC^*−*^ model (red). For the RC^+^ model, the fitted connectivity strengths, *w*_MN_ and *w*_RC_ *⊙ R*, between MNs and the RC population are not shown here. (2nd and 4th rows) Histograms of MU firing times after the TMS pulse in the RC^+^ model (black) and the RC^*−*^ model (red), stacked by TMS intensity (%MSO).

